# Interacting host modifier systems control *Wolbachia*-induced cytoplasmic incompatibility in a haplodiploid mite

**DOI:** 10.1101/2021.11.11.467240

**Authors:** Nicky Wybouw, Frederik Mortier, Dries Bonte

**Affiliations:** Terrestrial Ecology Unit, Department of Biology, Faculty of Sciences, Ghent University, Ghent, Belgium

**Keywords:** host modification, reproductive parasitism, genetic conflict, pest control

## Abstract

Many reproductive parasites such as *Wolbachia* spread within host populations by inducing cytoplasmic incompatibility (CI). CI occurs when parasite-modified sperm fertilizes uninfected eggs. In haplodiploid hosts, CI can lead to different phenotypes depending on whether the fertilized eggs die or develop into males. Genetic conflict theories predict the evolution of host modulation of CI, which in turn strongly influences the stability of reproductive parasitism. Yet, despite the ubiquity of CI-inducing parasites in nature, there is no conclusive evidence for strong intraspecific host modulation of CI strength and phenotype. Here, we tested for intraspecific host modulation of *Wolbachia*-induced CI in haplodiploid *Tetranychus* spider mites. Using a single CI-inducing *Wolbachia* variant and mitochondrion, a *Tetranychus urticae* nuclear panel was created that consisted of infected and cured near-isogenic lines. We performed a highly replicated age-synchronized full diallel cross comprised of incompatible and compatible control crosses. We uncovered host modifier systems that strongly suppress CI strength when carried by infected *T. urticae* males. Interspecific crosses showed that the male modifier systems suppress CI strength across species boundaries. We also observed a continuum of CI phenotypes in our crosses and identified strong intraspecific female modulation of CI phenotype when paired with a specific male genotype. Crosses established a recessive genetic basis for the maternal effect and were consistent with polygenic Mendelian inheritance. Our findings identify spermatogenesis as an important target of selection for host suppression of CI strength and underscore the importance of maternal genetic effects for the CI phenotype. Both mechanisms interacted with the genotype of the mating partner, revealing that intraspecific host modulation of CI strength and phenotype is underpinned by complex genetic architectures.

## Introduction

Endosymbiotic reproductive parasites facilitate their maternal transmission by manipulating host reproduction [1,2]. Cytoplasmic incompatibility (CI) is the most common reproductive manipulation and induces defects in paternal chromosome condensation and segregation in developing embryos when infected males mate with uninfected females [3]. In diploid hosts, these defects in paternal chromosome behavior result in embryonic mortality, whereas in hosts with a haplodiploid mode of reproduction (unfertilized haploid eggs develop into males, fertilized diploid eggs result in females), two different CI phenotypes can be distinguished. The chromosomal defects can give rise to viable haploid male offspring and is commonly referred to as Male Development CI (MD-CI) [4,5]. Alternatively, CI in haplodiploids can lead to aneuploidy in fertilized eggs and induce female mortality in incompatible crosses (Female Mortality CI, FM-CI) [4–6]. Reproductive parasites can induce a mix of both FM-CI and MD-CI in haplodiploid arthropods, suggesting that the phenotypic outcomes of incompatible crosses can lie along a continuum with FM-CI and MD-CI as the extremes [4–9]. CI is ablated when infected males mate with infected females and therefore provides a selective advantage to females that transmit the reproductive parasite.

CI is primarily associated with *Wolbachia*, but other bacteria such as *Rickettsiella, Cardinium*, and *Mesenet* can also induce CI in their arthropod host [3,10–13]. The genetic architecture of *Wolbachia*-mediated CI is a pair of syntenic genes (*cifA* and *cifB*) that are located in WO prophage regions of certain *Wolbachia* genomes [14–16]. Across different diploid and haplodiploid host systems, CI strength is highly variable and is typically associated with varying *Wolbachia* frequencies within host populations [17]. Although previous work has shown that natural genetic variation of *cif* operons can influence CI strength [18–20], *Wolbachia* genetic diversity does not sufficiently explain CI strength and phenotype variation. Interspecific transfer of *Wolbachia* and host species hybridization has revealed that host background can be a strong determining factor for CI strength and phenotype [21–26]. For example, *w*Mel *Wolbachia* induce weak CI in its native host *Drosophila melanogaster* [23,24], but cause strong CI in transinfected *Drosophila simulans* and *Aedes aegypti* [25,26]. In the haplodiploid wasp genus *Nasonia*, the *Wolbachia* strain determines CI strength, whereas interspecific crosses uncovered a role for host genotype in the expression of CI phenotype [8].

Interspecific modulation of CI is consistent with genetic conflict theories that predict the evolution of host modifier systems in populations that are polymorphic for *Wolbachia* infection [27]. However, intraspecific modulation of CI is not well documented, nor understood, in part due to variation caused by (asymmetrical) nuclear- or mitochondrial-associated incompatibilities [24,28–30]. Intraspecific variation in CI strength as well as phenotype has been observed for the haplodiploid spider mite *Tetranychus urticae* and the parasitoid wasp *Leptopilina heterotoma* [4–6,31], raising questions about the ability of haplodiploid hosts to control the different features of CI. Despite the ubiquity of CI in the natural world, the mechanistic underpinnings of host modulation are not known but can be characterized along several axes. Host modulation mechanisms can be expressed in males, females, and embryos of incompatible crosses. In males, these mechanisms limit *Wolbachia*-induced aberrations of spermatogenesis to produce normal, healthy sperm. Females can modify CI through maternal effects that operate during oogenesis and embryogenesis. For instance, maternally supplied histones are vital for the formation of the male pronucleus and display a delayed deposition in embryos from incompatible crosses, findings that identify this process as a potential target for host selection [32,33]. Finally, the embryo can also modulate CI throughout its development. Although CI is commonly associated with defects in the early mitotic divisions [3], cytological assays have uncovered additional CI phenotypes in later stages of embryonic development, such as regional mitotic failures [15]. Moreover, *Wolbachia*-induced CI can also result in postembryonic mortality [7], dynamics that could allow for further selection for modifier systems in the offspring of incompatible crosses. Whether host modifier systems control different features of CI or combine additively or synergistically is unknown, and are pertinent questions for our understanding of reproductive parasitism and the development of effective CI-based pest management. Here, reproductive parasites are released into genetically diverse pest populations across the globe and reduce population growth by inducing strong CI. Segregating modifier systems in wild pest populations can severely threaten the long-term stability and effectiveness of CI-based pest control [34,35].

To study the mechanistic underpinnings of host modulation in a haplodiploid arthropod, we created a nuclear panel of the spider mite *T. urticae* comprised of different nuclear backgrounds, a single mitochondrion, and a single CI-inducing *Wolbachia* variant. We performed a highly replicated full diallel cross and quantified variation in CI strength and phenotype using Bayesian inference and corrected indexes that control for nuclear and temporal effects. CI strength varied from very weak to complete and was strongly determined by the male genotype. CI phenotypes ranged along a continuum and were determined by the female and male genotype as well as their interaction. Interspecific crosses were performed to study the control of male modifier systems when paired with a female genotype of a different *Tetranychus* species. Genetic crosses uncovered insights into the genetic architecture underlying intraspecific female modulation of the CI phenotype. Together, our findings reveal multiple mechanisms of host modulation of CI strength and phenotype within a single haplodiploid arthropod species.

## Materials and Methods

### Creation and characterization of the *Tetranychus* nuclear panel

A teleiochrysalid (virgin) female was collected from five *Tetranychus* field populations (Bch, Beis, Scp-*w*, Stt, and Temp) and from a laboratory population (LonX) that was derived from the London reference strain [36] (Table S1). To ensure near-isogenic nuclear backgrounds, lines were created by three sequential rounds of mother-son crosses. Genomic DNA was extracted from a pool of 20 adult females with a Quick-DNA Universal kit (BaseClear, the Netherlands). A fragment of the mitochondrial *COI* gene was sequenced for the Scp-*w* and Bch lines (Sanger sequencing, MACROGEN Europe B.V.) (Table S2). Infection of the six *Tetranychus* lines with the reproductive manipulators *Wolbachia, Rickettsia, Cardinium*, and *Spiroplasma* was tested using diagnostic PCR assays. PCR assays were performed using DreamTaq DNA Polymerase (Life Technologies Europe B.V.) in a 50 μl reaction mixture. PCR conditions are described in Table S2. A *Wolbachia*-infected *Myrmica scabrinodis* worker and laboratory populations of *T. urticae* that were infected with *Wolbachia*, *Rickettsia, Cardinium*, and *Spiroplasma* were used as positive controls. The diagnostic PCR assays showed that none of the lines carried *Rickettsia, Cardinium*, or *Spiroplasma* and that only Scp-*w* was infected with *Wolbachia*. The *Wolbachia* variant of Scp-*w* was further characterized by multilocus sequence typing (Sanger sequencing, MACROGEN Europe B.V.) (Table S2) [37]. We transferred the *Wolbachia* variant into four other near-isogenic backgrounds by paternal introgression, creating Beis-*w*, LonX-*w*, Stt-*w*, and Temp-*w*. For each near-isogenic nuclear background, 20 *Wolbachia*-infected Scp-*w* females were crossed to 15 uninfected males, and 25 infected female offspring were backcrossed to 15 males of the uninfected genotype for an additional six generations (figure 1A). Introgressive transfer of *Wolbachia* into the Bch nuclear background failed since F_1_ females did not oviposit. Hybrid sterility is a pervasive post-zygotic reproductive barrier between tetranychid species [38,39]. We morphologically classified the Bch line (electronic supplementary material). The fidelity of *Wolbachia* maternal transmission was tested in the five infected nuclear backgrounds of *T. urticae*. DNA was extracted from individual adult females and males (electronic supplementary material) and the diagnostic PCR assays were designed as described above (Table S2). The total numbers of tested mites are listed in Table S3. The Beis-*w*, LonX-*w*, Scp-*w*, Stt-*w*, and Temp-*w* lines were cured of *Wolbachia* infection by antibiotic treatment (electronic supplementary material). After antibiotic curing, the cured lines Beis-c, LonX-c, Scp-c, Stt-c, and Temp-c were maintained on detached bean leaves for at least four generations before *Wolbachia*-mediated incompatibilities were phenotyped. The 11 lines of the mite panel were maintained by serial passage on detached bean leaves at a census size of ~250 mites at 24°C, 60%RH, and a 16:8 light:dark photoperiod.

**Figure 1.**
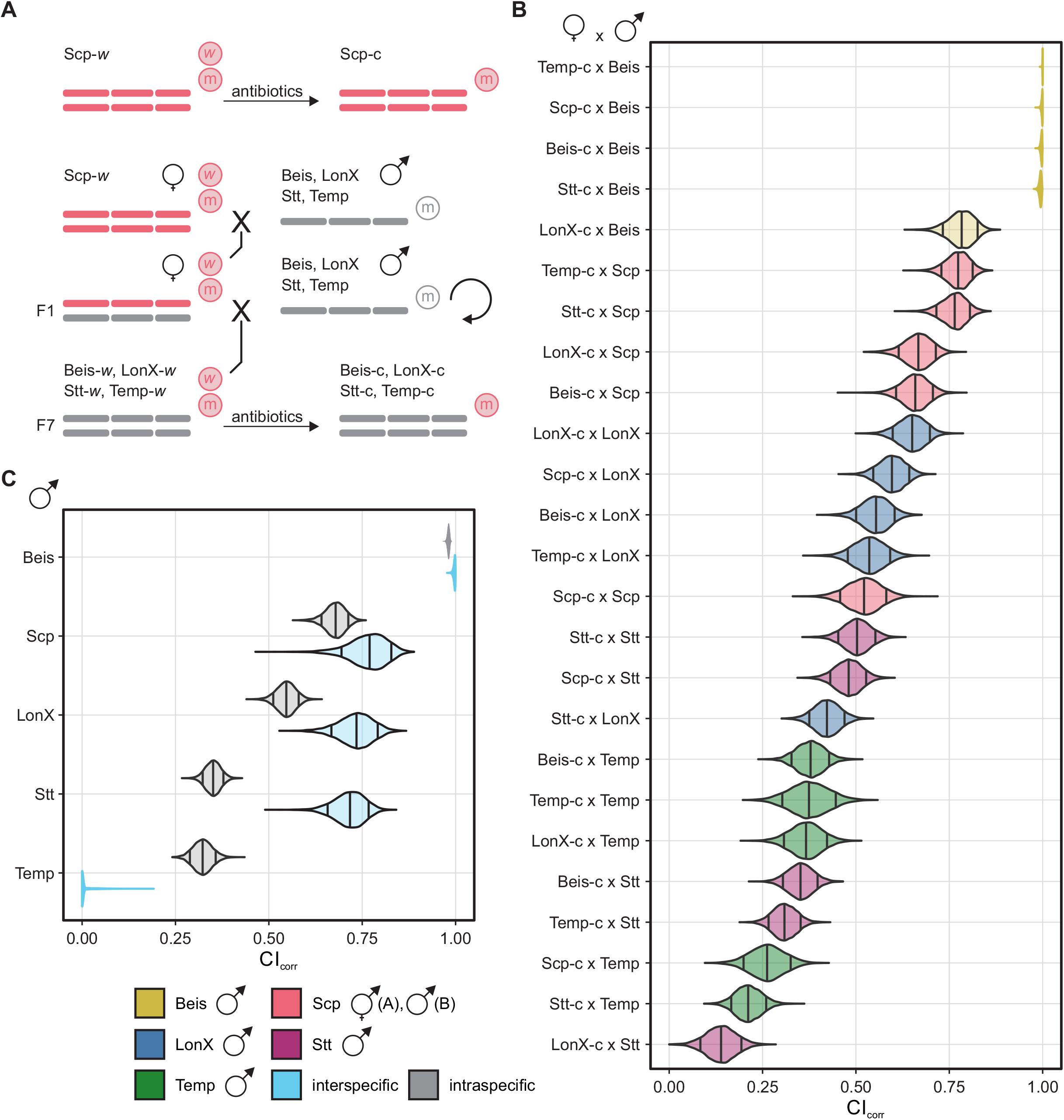
Host modifier systems suppress CI strength in *Tetranychus* spider mites. **(A)** The experimental design that created the *T. urticae* nuclear panel. *Wolbachia* was transferred from Scp-*w* into the Beis, LonX, Stt, and Temp nuclear backgrounds by paternal introgression, creating Beis-*w*, LonX-*w*, Stt-*w*, and Temp-*w*. Infected lines were cured of *Wolbachia* by antibiotic treatment, creating Beis-c, LonX-c, Scp-c, Stt-c, and Temp-c. The *Wolbachia* variant and mitochondrion of Scp-*w* are indicated by encircled ‘*w*’ and ‘m’ symbols in red font. **(B)** Intraspecific CI strength variation within the full diallel cross design. Crosses are ordered according to decreasing CI strength. Violin plots are colour coded based on the male genotype (see bottom left). **(C)** Interspecific CI strength variation using five *T. urticae* male genotypes and uninfected Bch females. Bch is a distinct *Tetranychus* species. Crosses are ordered according to decreasing intraspecific CI strength. Violin plots are colour coded based on whether the cross was intraspecific or interspecific (see bottom left). For (B) and (C), CI strength was estimated using the CI_corr_ index that controls for variation caused by nuclear and temporal effects. Each violin plot represents the estimated average of CI_corr_ for that cross and indicates the 0.09, 0.50 and 0.91 percentiles.

### Host modulation of *Wolbachia*-induced CI

Host modifier systems were identified and characterized in our *T. urticae* panel by performing incompatible (uninfected females to infected males) and compatible (uninfected females to uninfected males) intraspecific crosses in a full factorial diallel cross design. This experimental design generated a total of 50 intraspecific cross types; 25 incompatible and 25 compatible control cross types. The control of *T. urticae* modifier systems in males of interspecific CI was characterized by crossing *T. urticae* males of the five near-isogenic nuclear backgrounds to uninfected Bch females, generating an additional set of five incompatible and five compatible control cross types. To provide further validation that Bch represents a distinct species from *T. urticae*, five F_1_ female teleiochrysalids were isolated per cross type (if present) and allowed to oviposit on 16 cm^2^ leaf discs until death. A third set of crosses tested the ability of infected females of Scp-*w* and Beis-*w* to rescue CI and consisted of rescue (infected females to infected males) and control (infected females to uninfected males) crosses. Age cohorts of the mite lines were created by allowing 50 mated females to oviposit for 24 hours on detached bean leaves. Each cross type consisted of eight to eleven replicates except for the rescue and respective control crosses where five replicates were established per cross type (electronic supplementary material). For each replicate, five female teleiochrysalids were paired with four one- to three-day old adult males on a 16 cm^2^ leaf disc. After five days, mites were discarded and eggs were counted. During development, adult male and female offspring were isolated and counted.

### Genetic basis of female modulation of CI phenotype

We uncovered the mode of inheritance of female modulation of CI phenotype using Scp-c and LonX-c females that express distinct CI phenotypes when crossed to Scp-*w* males (figure 2A). Heterozygous F_1_ females were produced by reciprocal crosses between LonX-c and Scp-c. In each cross, 25 virgin females were paired with 20 adult males on a 16 cm^2^ leaf disc and allowed to mate and oviposit. During F_1_ development, female teleiochrysalids were isolated for the incompatible and compatible control crosses. Adult Scp-*w* and Scp-c males were obtained from synchronized age cohorts as previously described. Cross types consisted of seven to eight replicates, with five females and four males per replicate. After five days, mites were discarded and eggs were counted. During development, adult male and female offspring were isolated and counted.

**Figure 2.**
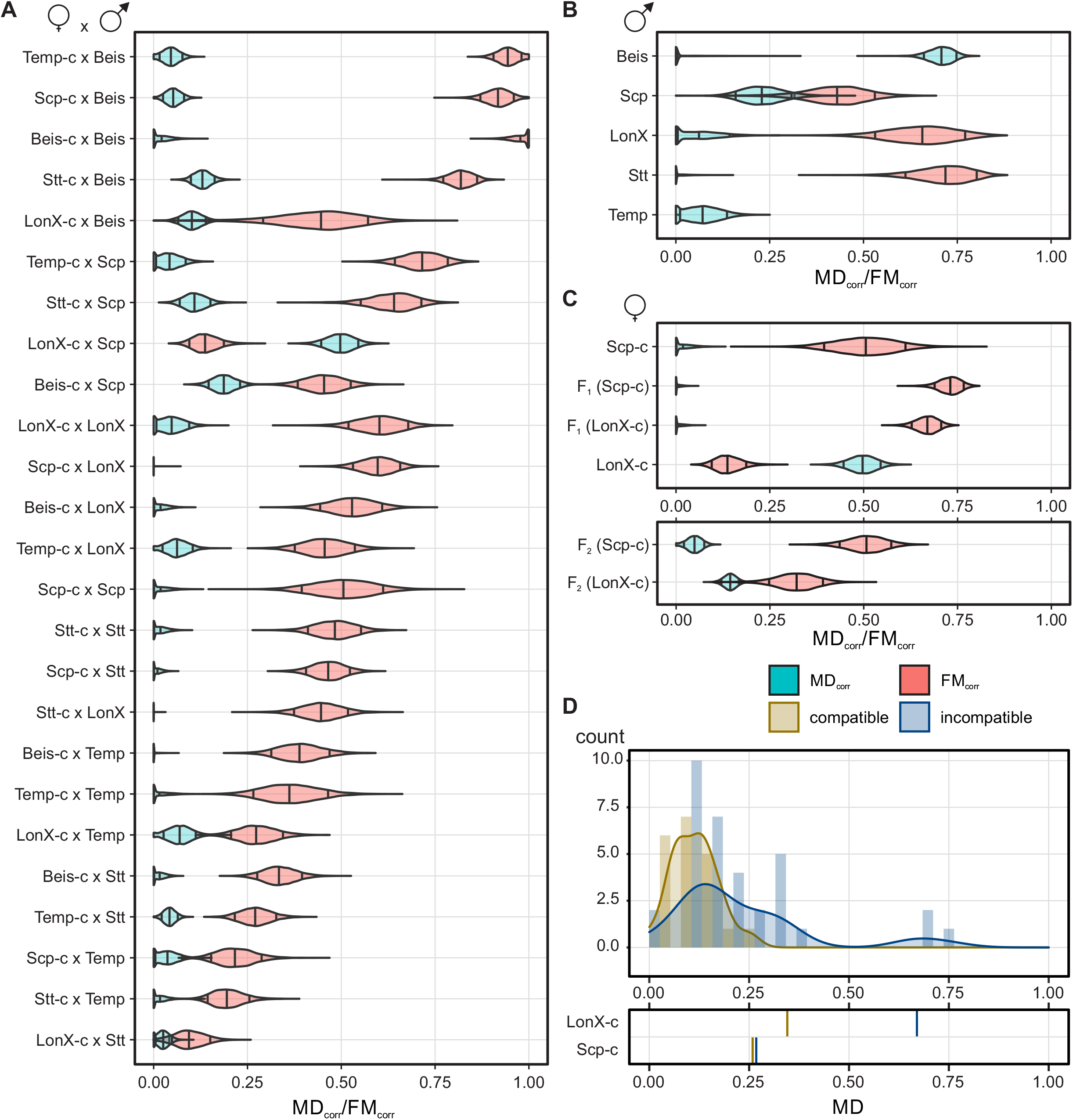
Host modulation of CI phenotype in *Tetranychus* spider mites. **(A)** Intraspecific CI phenotype variation within the full diallel cross design. Crosses are ordered according to decreasing CI strength. **(B)** Interspecific CI phenotype variation using five *T. urticae* male genotypes and uninfected Bch females. Bch is a distinct *Tetranychus* species. Crosses are ordered according to decreasing intraspecific CI strength. **(C)** Inheritance of the maternal genetic effect that contributes to intraspecific MD-CI variation. For the heterozygous F_1_ and recombinant F_2_ females, the genotype between brackets represents the original maternal genotype. All uninfected females were crossed to Scp-*w* and Scp-c males. MD_corr_ and FM_corr_ estimates of LonX-c and Scp-c are identical to those of panel A. **(D)** Distribution of MD in the incompatible and compatible crosses of recombinant F_2_ females and Scp-*w* and Scp-c males, respectively. Cross compatibility is colour coded (see middle right). The average MD values for LonX-c and Scp-c are shown in the bottom plot. For all panels, violin plots of MD_corr_ and FM_corr_ display a blue and red background, respectively (see middle right). Violin plots represent the estimated averages of MD_corr_ and FM_corr_ for each cross and indicate the 0.09, 0.50 and 0.91 percentiles.

To obtain recombinant F_2_ females, 25 virgin F_1_ females were backcrossed to 20 LonX-c males. Individual female F_2_ teleiochrysalids were isolated, paired with a single one-to three-day old adult Scp-*w* or Scp-c male, and allowed to oviposit for seven days. Control cross types consisted of five to 21 replicates, whereas the incompatible cross types included 22 to 34 replicates. During development, adult male and female offspring were isolated and counted.

### Statistical analysis of CI strength and phenotype

CI strength and phenotype were analyzed with Bayesian inference using the brms package (version 2.12.0) in R (version 3.6.3) [40,41]. Statistical models were run using Hamiltonian Monte Carlo (HMC) that implemented two chains with each 5,000 iterations from which 2,000 were warmup. We chose error distributions with ‘biggest entropy’; the distribution with the least amount of assumptions that is consistent with the constraint of the outcome variable from first principles [42]. We modelled the proportion of adult female offspring over total number of eggs (F), proportion of adult male offspring over total number of eggs (MD), and the proportion of eggs that failed to generate adult mites over total number of eggs that did not generate adult males (FM) as responses with a binomial error distribution. These will be used to calculate informative CI, MD-CI and FM-CI metrics, respectively (see further). Priors were used that are weakly regularizing by choosing prior distributions that are significantly wider than the parameter values that would be reasonable to expect (electronic supplementary material). We evaluated the performance of every fitted model based on standardized procedures by checking mixing and stationarity in the trace plots and by checking the effective sample size and *Ȓ* statistic for each parameter [42].

We estimated F, MD, and FM using full models. For intraspecific crosses, F, MD, and FM were modelled with effects from the *Wolbachia*-infection state in *T. urticae* males, male genotype, female genotype, and all their interactions. For interspecific crosses, F, MD, and FM were modelled with effects from the *Wolbachia*-infection state in *T. urticae* males, male genotype, whether the cross was inter-or intraspecific, and all their interactions. Here, we assumed no effect of the *T. urticae* female genotype based on our model comparisons of the intraspecific crosses (see further). For all analyses above, the days at which the different cross types were initiated were included as variable intercepts. To study CI rescue, we estimated F using the full model that incorporates all effects and their interactions. To study the inheritance of female modulation, F, MD, and FM were modelled with the effects from the *Wolbachia*-infection state in males and the female genotype. For CI rescue and female modulation, the experimental design did not allow to estimate the variable day effect.

To control for variation in F, MD, and FM that is not related to *Wolbachia*-induced CI (caused by nuclear and temporal effects), we used corrected indexes. Using the model estimates of F, the corrected CI strength (CI_corr_) for each cross type was calculated from the posterior distribution of the full generalized linear model:

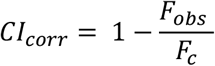

where F_obs_ and F_c_ are the estimated F values in the incompatible and respective compatible crosses, respectively [7,31]. The corrected MD-CI phenotype (MD_corr_) was calculated using the estimated MD values;

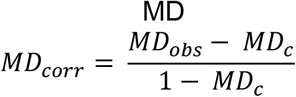

where MD_obs_ and MD_c_ are the estimated MD values in the incompatible and respective compatible crosses, respectively [25,39,43]. The corrected FM-CI phenotype (FM_corr_) was calculated using the estimated FM values;

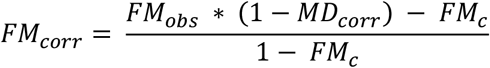

where FM_obs_ and FM_c_ are the estimated FM values in the incompatible and respective compatible crosses, respectively (based on [25,39,43]).

To understand the importance of male and female genotype and their interaction for intraspecific CI strength and phenotype, we compared four variations of the full models that consistently assumed an effect of the *Wolbachia*-infection state in males but differed in the other fixed explanatory variables (male genotype, female genotype and their interactions) (electronic supplementary material). Models were compared using the Widely-Applicable information criterion (WAIC). To quantify the relative impact of the different explanatory variables, we compared the finite-population standard deviation of estimated coefficients for different explanatory variables and interactions in adjusted full models (electronic supplementary material). These adjusted full models estimated all effects as modelled variable effects instead of as fixed to improve our ability to interpret coefficients and their standard deviation accurately in a model with interactions. The standard deviation of the coefficients of the multiple model levels were visualized on the log-odds scale and compared to the standard deviation of the model residuals as a reference of unexplained variation.

## Results

We created a *T. urticae* panel comprised of five near-isogenic nuclear backgrounds that shared a single mitochondrion and were either infected with a single CI-inducing *Wolbachia* variant (Beis-*w*, LonX-*w*, Scp-*w*, Stt-*w*, and Temp-*w*) or were cured of the infection (Beis-c, LonX-c, Scp-c, Stt-c, and Temp-c). A complete maternal transmission of *Wolbachia* was observed in all five infected *T. urticae* nuclear backgrounds (Table S3). All intraspecific compatible crosses produced F_1_ females, demonstrating fertilization across all near-isogenic lines (electronic supplementary material). Morphological classification showed that the Bch line is distinct from *T. urticae* (electronic supplementary material) and compatible control crosses of *T. urticae* males of all near-isogenic lines and uninfected Bch females produced hybrid F_1_ females. Virgin F_1_ females did not oviposit across our cross types (n= five females per cross type, if present), a post-zygotic reproductive barrier that is commonly observed in interspecific crosses with *Tetranychus* mites [38,39].

### Male modifier systems cause weak CI in *Tetranychus*

Using the model estimates of the full model (figure S1), we calculated the corrected CI strength (CI_corr_) across the intraspecific cross types, controlling for nuclear and temporal effects (figure 1B). Intraspecific CI_corr_ varied greatly and ranged from complete to very weak. A model comparison was performed among models that differed in fixed explanatory variables to study the factors explaining intraspecific CI strength variation (figure S2). These model comparisons revealed that the male genotype had the largest impact on model predictability, with its interaction with the female genotype as an important additional effect (figure S2). Variance analysis confirmed that the male genotype greatly determined intraspecific CI strength with an additional substantial impact of the male-female genotype interaction (figure S3). In contrast, the coefficients of all other model levels exhibited markedly lower levels of variation (figure S3). These analyses indicate that (some) *T. urticae* male genotypes carry modifier systems that strongly modulate intraspecific CI strength.

*Wolbachia*-infected Beis-*w* males induced complete (or near-complete) CI_corr_ when crossed to four uninfected female genotypes (Beis-c, Scp-c, Stt-c, and Temp-c). In contrast, all other genetic crosses revealed reduced levels of CI_corr_. Weakest CI_corr_ was observed in the crosses with infected Stt-*w* and Temp-*w* males, whereas infected Scp-*w* and LonX-*w* males induced intermediate levels of CI_corr_ (figure 1B). The level of interaction of the male and female genotype on CI_corr_ varied across the intraspecific crosses (figure 1B and figure S4). The strongest interaction was observed when infected Beis-*w* males were crossed to uninfected LonX-c females, resulting in a CI_corr_ of ~75% (figure 1B and figure S4). Together, these findings suggest that the LonX, Scp, Stt, and Temp genotypes carry nuclear modifier systems that are expressed in males and strongly reduce intraspecific CI strength. The ability of infected females to rescue intraspecific CI was confirmed for the Scp and Beis genotypes using replicated age-synchronized rescue (infected females to infected males) and control cross types (infected females to uninfected males) (figure S5).

The ability of Bch and *T. urticae* to hybridize allowed us to test the strength of the modifier systems in *T. urticae* males across species boundaries (figure 1C). Here, we found that infected Beis-*w* males continued to induce complete CI_corr_, whereas crosses with the other *T. urticae* male genotypes resulted in lower levels of CI_corr_ (figure 1C). Infected Temp-*w* males did not induce CI_corr_ when crossed to uninfected Bch females, indicating a complete suppression of CI by the host within this cross.

### Male modifier systems suppress CI by controlling female mortality

The total CI strength in haplodiploids is the combined effect of MD-CI and FM-CI where the incompatibility is expressed as an excess of haploid male offspring or an increased mortality of female offspring, respectively. Using the model estimates of the full models, we calculated the corrected MD-CI and FM-CI indexes (MD_corr_ and FM_corr_, respectively) (figure S6 and figure S7). We observed a continuum for both CI phenotypes within *T. urticae*, but FM_corr_ variation was considerably larger (figure 2A). The variation of MD_corr_ and FM_corr_ across the intraspecific crosses was examined by running several statistical models with the *Wolbachia*-infection state in males as a consistent explanatory variable. Model comparisons for MD-CI indicated that male and female genotype and their interaction contributed equally to model predictability (figure S8) but accounted for a relatively low amount of variation (figure S9). In contrast, model comparisons and variance analysis uncovered that the male genotype and, to a lesser extent, its interaction with the female genotype were important determinants for FM-CI, showing a correlation between CI strength and FM-CI in our data (figure S8 and figure S9). Correlation plots confirmed that FM_corr_ and CI_corr_ were tightly coupled, whereas patterns with MD_corr_ were inconsistent (figure S10). Together, this suggests that the control of male modifier systems of intraspecific CI strength was mainly regulated by changes in the mortality rate of female offspring (figure 2A).

Crossing LonX-*w*, Scp-*w*, and Stt-*w* males to uninfected Bch females resulted in interspecific CI that was mainly determined by FM_corr_ (figure 2B). In contrast, strong MD_corr_ was observed when Beis-*w* males were crossed to Bch (figure 2B), revealing an interaction effect between the Beis and Bch genotypes for the expression of interspecific MD-CI.

### A recessive maternal genetic effect contributes to intraspecific MD-CI variation

The LonX-c (♀) x Scp-*w* (♂) cross was the only intraspecific incompatible cross where MD_corr_ (~50%) exceeded FM_corr_ (~12.5%) (figure 2A and figure S10). Crossing Scp-*w* males to other uninfected female genotypes caused no or very weak intraspecific MD_corr_ (figure 2A), suggesting that the relatively strong MD_corr_ resulted from an interaction between a maternal genetic feature of LonX with the Scp genotype. To gain further insight into female modulation of CI, we produced heterozygous F_1_ females by reciprocal crosses between LonX-c and Scp-c, and performed replicated incompatible and compatible crosses with Scp males (figure 2C). CI strength was stable between parents and F_1_ offspring with the incompatible crosses of both sets of F_1_ females and Scp-*w* males resulting in ~70% CI_corr_ (figure S11). Crossing heterozygous F_1_ females to Scp-*w* males did not cause MD_corr_, establishing a recessive genetic basis for the LonX maternal effect (figure 2C). We subsequently backcrossed F_1_ females to LonX-c males, and performed replicated incompatible and compatible crosses using a single recombinant F_2_ female per cross. CI_corr_ remained stable and was ~70% (figure S11). We observed a ~5% and ~15% MD_corr_ after crossing recombinant F_2_ females to Scp-*w* (figure 2C). We noted a clear change in the distribution of MD using recombinant F_2_ females with LonX-c as the original maternal line (figure 2D). After mating with Scp-*w* males, 3 out of 34 (8.8%) recombinant F_2_ females produced a brood with an MD value that exceeded the average MD of LonX-c (figure 2D), patterns that are consistent with a polygenic basis.

## Discussion

Intraspecific host modulation of parasite-induced CI is predicted to influence the evolutionary trajectory of host-parasite interactions and, by suppressing CI strength, the long-term effectiveness of CI-based pest management [27,34,44]. Here, we identified strong host modulation by the spider mite *T. urticae* of *Wolbachia*-mediated CI. We collected convincing evidence that a nuclear modifier system, or systems, segregates in *T. urticae* that strongly suppresses CI strength when carried by *Wolbachia*-infected males. The strongest suppression of CI was observed in infected Stt-*w* and Temp-*w* males (intraspecific CI strength dropped as low as ~15% CI_corr_). Crosses of *T. urticae* males to uninfected Bch females showed that the male modifier systems also strongly suppress CI strength across species boundaries. Complete (or near-complete) CI was only observed in the genetic crosses with infected Beis males, suggesting that only the Beis genotype lacks male suppressors within our *T. urticae* panel. Despite the ubiquity of CI-inducing parasites, evidence for intraspecific host suppression of CI strength is rarely reported. Genetic work on *Drosophila* and *Nasonia* has gathered some evidence of modest intraspecific host modulation of CI strength [24,28,45]. However, these results remain largely inconclusive due to a lack of control for (asymmetrical) nuclear-or mitochondrial-associated incompatibilities and line homozygosity. In *D. simulans, Aedes aegypti*, and *Culex pipiens*, no intraspecific genetic variation has (yet) been observed that modulates CI strength despite intensive sampling efforts [29,30,44,46,47]. These findings are in sharp contrast with our study where we uncovered host suppression in four out of five isogenic lines. The maternal transmission of *Wolbachia* in tetranychid mites is sensitive to increased temperatures and is likely to be imperfect under natural conditions, preventing *Wolbachia* from reaching fixation and resulting in a persistent expression of CI [48–50]. Our data therefore support models that predict the evolution of host modifiers in systems with strong CI and imperfect maternal transmission [2,27].

In contrast to CI, host nuclear suppressors that act against male-killing, a less common parasite-induced manipulation, have been described in a wide range of arthropod species, including fruit flies, butterflies, and spiders [2,51–53]. CI and male-killing exert different selection pressures on the arthropod host [2]. Male-killing reduces the fitness of infected females that transmit the parasite, whereas CI has deleterious effects for uninfected females. Moreover, in contrast to male-killing, the fitness cost of CI is ablated in populations that are fixed for the reproductive parasite [2,27]. These divergent fitness penalties could explain the observed discrepancy in host modulation. Additionally, parasite-mediated male-killing of infected males is achieved by manipulating the dosage compensation system that regulates sex differentiation [54,55] and these pathways may be intrinsically more polymorphic and display greater standing genetic variation within host populations compared to those that are influenced by parasite-mediated CI.

The mechanism by which infected males are conditionally (partially) sterilized by *Wolbachia* is not well understood [3,16,56], limiting our ability to unravel the mechanistic basis of host modulation. The host can develop toxicodynamic resistance to CI by changes in the target-sites of *Wolbachia* Cif proteins (coined as the defensive model in [3]). As *Wolbachia* density has been observed to positively covary with CI strength and maternal transmission efficiency [3,57], CI can be overcome by genetic variants in the host that dysregulate *Wolbachia* density, a mechanism that can be viewed as toxicokinetic resistance (coined as the offensive model in [3]). In *Nasonia* wasps, host genetics of infected females contribute to variation in the maternal transmission of *Wolbachia* to the progeny [58]. In our study, maternal transmission of *Wolbachia* appeared complete in all infected genetic backgrounds, including the Stt and Temp genotypes that exhibited the strongest male suppression of CI. The interaction of the male modifier systems with the uninfected female genotype was an important determinant for CI suppression, suggesting that variation in *Wolbachia* density in infected males does not (fully) explain CI strength variation across our diallel cross design. In *Wolbachia*-infected wasps and fruit flies, CI strength is coupled with *Wolbachia cif* gene expression [15,59], and host modulation of *cif* transcription could underlie the variation in CI strength observed in our study. *Cif* genes are divided into a minimum of five phylogenetic clades and the encoded proteins exhibit extensive variation in domain structure [19]. Although all CifB proteins have a dimer of PD-(D/E)XK nuclease domains, the diversity of additional functional domains indicate that CI may be manifested by different biochemical mechanisms across different *Wolbachia* variants, a hypothesis that finds some support in previous work [3,16,19]. Unfortunately, the *cif* repertoire of *Wolbachia* that infect *T. urticae* is not identified. Previous attempts to amplify and sequence *cifA* and *cifB* genes of *Wolbachia* in *T. urticae* using reference primers failed [43], indicating high gene sequence divergence from previously characterized *cif* genes. Multilocus sequence typing showed that the CI-inducing *Wolbachia* variant of the current study belonged to supergroup B, a clustering that is consistent with all previously characterized variants in *T. urticae* [43,60]*. Wolbachia* infection of *Tetranychus* mites is characterized by an apparent high strain diversity [60], raising the question of how the host modifier systems of our mite panel would interact with *Wolbachia* variants that carry divergent *cif* repertoires. Although speculative, the high prevalence of weak and intermediate CI in *T. urticae* populations across the globe could be (partially) caused by different male suppressors that segregate at high frequencies [4,43,61]. In *Drosophila teissieri*, genetic crosses suggest that CI strength could be determined by an interaction between the *Wolbachia* variant and host genotype, but formal evidence awaits [28]. In *D. melanogaster*, suppressors of parasite-induced male-killing are located on different chromosomes and induce distinct suppressor phenotypes, indicative of independent origins of host suppression [52]. Further work is required to fully understand the mechanisms of male suppression of CI in our *T. urticae* genotypes.

We also gathered evidence of strong intraspecific female modulation of CI phenotype when paired with the Scp male genotype. In the wasp genus *Nasonia*, differences in CI phenotype across species is also (partially) attributed to female modulation [8]. Consistent with the *Nasonia* system, the maternal genetic effect that contributes to MD-CI in LonX is recessive [8]. Although the results of the backcross experiments appear consistent with polygenic Mendelian inheritance, the number of loci involved, their additivity and effect size remains unresolved. Multiple mechanisms can underpin female modulation of MD-CI. During embryogenesis, the maternal genetic effect could contribute to the complete elimination of the paternal chromosomes, giving rise to viable haploid male offspring. Using high-copy library plasmids in yeast and a protein-interaction screen in *Drosophila*, Beckmann *et al*. collected evidence that shows that a particular type of *Wolbachia* CifB interacts with the maternally deposited proteins karyopherin-α and P32, and identified protamine-histone exchange and nuclear-protein import as target pathways [62]. As nuclear transport has previously been associated with genetic conflict [63,64], it is tempting to speculate that these pathways underpin the observed maternal genetic effect of this study. Alternatively, the maternal genetic effect could result in (partial) fertilization failure by physiological changes within the female reproductive tissue. Future experiments that implement unbiased forward genetics are needed to uncover the genetic architecture of female modulation of CI phenotype.

To conclude, we identified mechanisms of intraspecific host modulation that suppress CI strength by male modifier systems and modify CI phenotype by maternal genetic effects. As both mechanisms interacted with the genotype of the mating partner, we show that different complex genetic architectures underlie intraspecific host modulation of parasite-induced CI.

## Supporting information

electronic supplementary material

electronic supplementary material

electronic supplementary material

electronic supplementary material

electronic supplementary material

## Author Contributions

NW conceived and designed experiments. NW and FM performed experiments. NW, FM, and DB analyzed data. NW wrote the manuscript with input from FM and DB.

## Acknowledgements

We are indebted to Tadek and Helena Wybouw for their assistance during the field collections. We thank Philippe Auger for performing the morphological classification of the *Tetranychus* Bch line. NW was supported by a BOF post-doctoral fellowship (Ghent University, 01P03420) and by a Research Foundation - Flanders (FWO) Research Grant (1513719N). The authors declare no conflicts of interest.

## Data Accessibility

The multilocus sequence type data of the CI-inducing *Wolbachia* variant are accessible at NCBI (OK669102-OK669106). The *COI* sequence data of the Bch and Scp-*w* line are accessible at NCBI (OL333561 and OL333562, respectively). Raw count data and scripts will be publicly available upon acceptance, or upon request.

