## Supplementary material for "Interacting host modifier systems control *Wolbachia*-induced cytoplasmic incompatibility in a haplodiploid mite": electronic supplementary material

### Supplementary figures

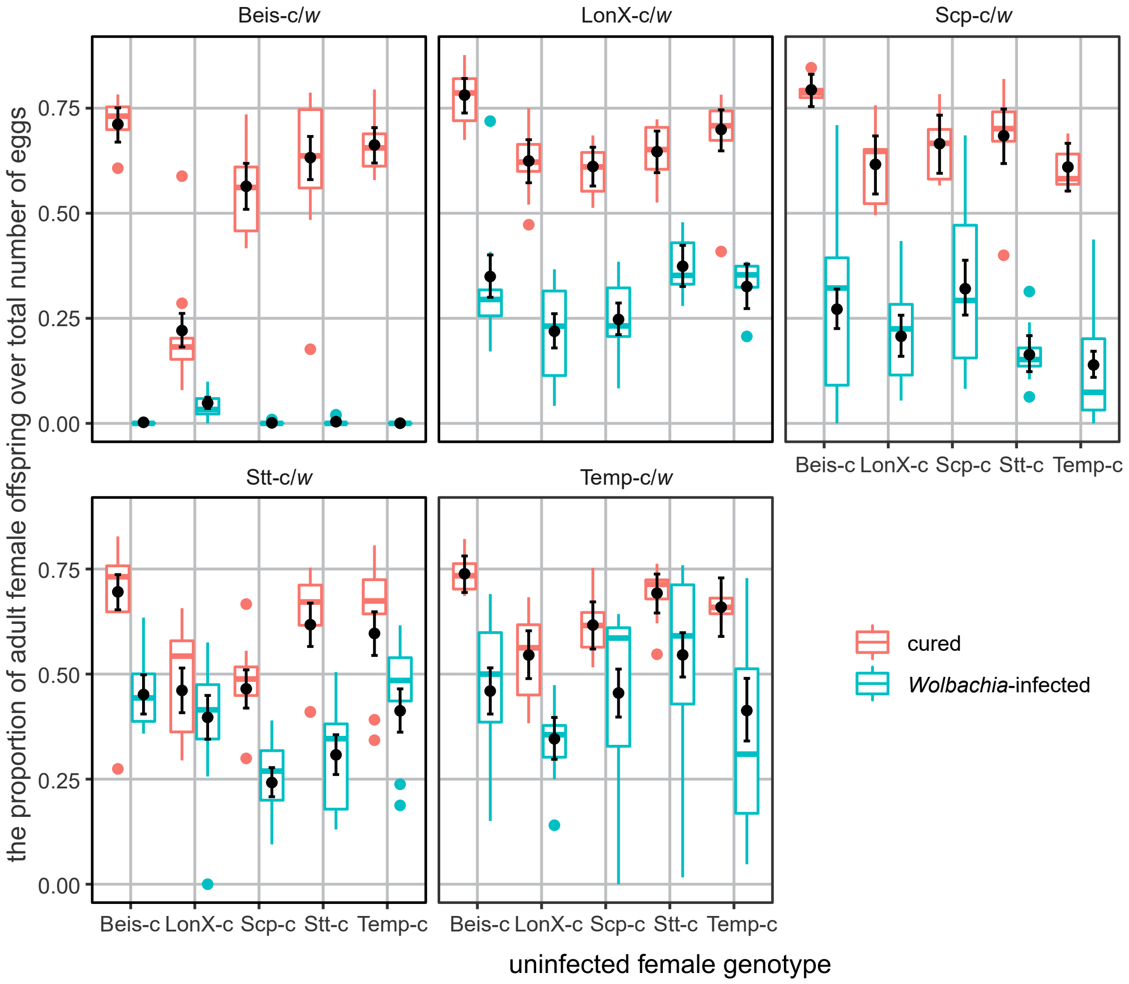


**Figure S1. Observed and estimated proportion of adult females over total number of eggs across the intraspecific cross types.** The panels are organized based on the male genotype. The infection state of the males is colour coded (see bottom right).The boxplots depict the raw data whereas the dot and whiskers depict the mean model estimates and the 0.09-0.91 likelihood intervals respectively from the HMC model (model (1)) without the variable intercept for measuring day (electronic supplementary material).

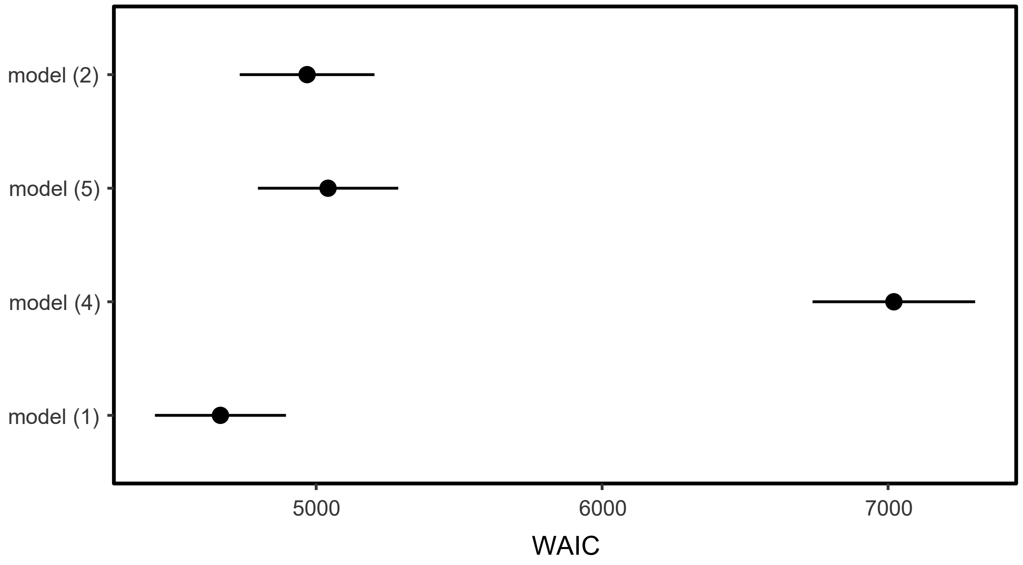


**Figure S2. Model comparisons for the analysis of intraspecific CI strength variation.** The WAIC scores and their standard errors are depicted for four models that were run to analyze intraspecific CI strength. The metrics of four models are depicted; a full model that includes the effects of male and female genotype with their interaction as explanatory variables (model (1)), a model that incorporates the effects of male and female genotype without their interaction (model (2)), a model that only incorporates the effects of female genotype (model (4)) or male genotype (model (5)). All models contained the effect of the *Wolbachia*-infection state of males and its interaction(s) with the other variable(s) and a random intercept for the days on which the cross types were initiated. Model comparisons uncovered that model (1) performed the best in terms of out-of-sample prediction. Model (5) performed only slightly worse than model (1), whereas model (4) performed the worst. Model (2) performed equally well as model (5), showing that information of the female genotype did not improve model performance when the male genotype was already taken into account but the reverse improved the model greatly. Detailed model descriptions can be found in the electronic supplementary material.

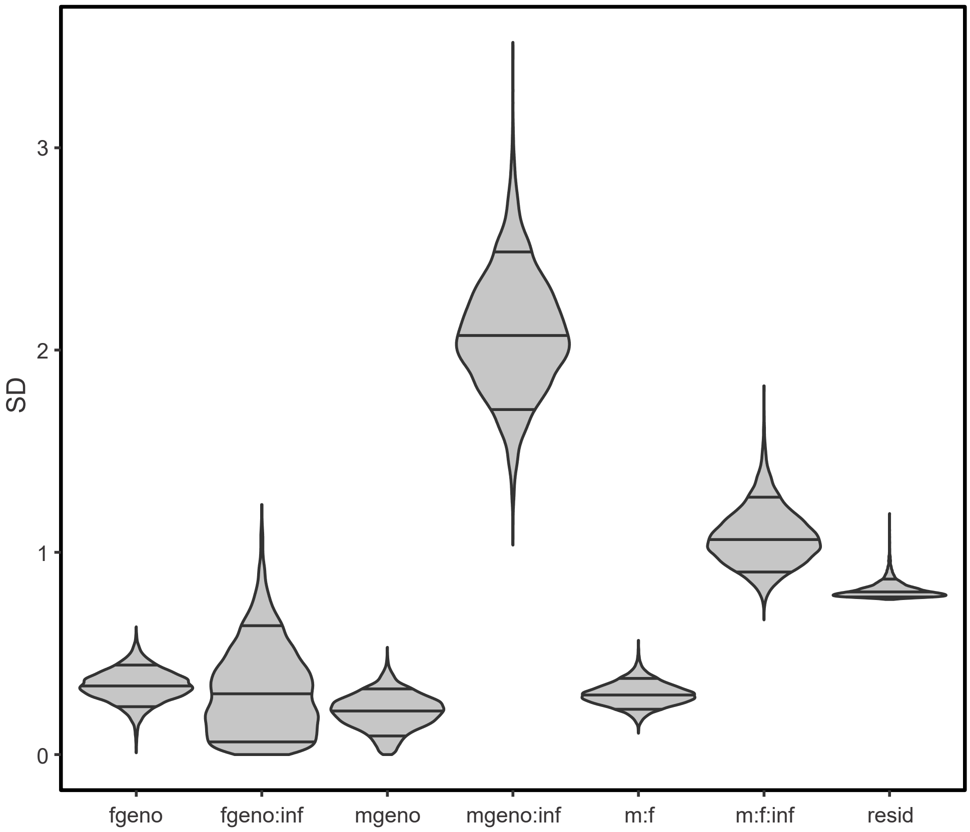


**Figure S3. Estimated finite-population standard deviation (SD) of coefficients of the levels of model (3) for intraspecific CI strength to quantify the relative impact of the explanatory variables.** For the analysis of intraspecific CI strength variation, the levels of the female genotype (fgeno), male genotype (mgeno), their interaction (f:m), and all the interactions with the *Wolbachia*-infection state in males (inf) were included. The standard deviation of the log-odds residuals of the model was included as a reference for the unexplained variance. The strongest interaction with the *Wolbachia*-infection state in males was the male genotype, while the second strongest interaction was the three-way effect with male and female genotype. The SD distributions show the 0.09, 0.5, and 0.91 percentiles. The detailed description of model (3) can be found in the electronic supplementary material.

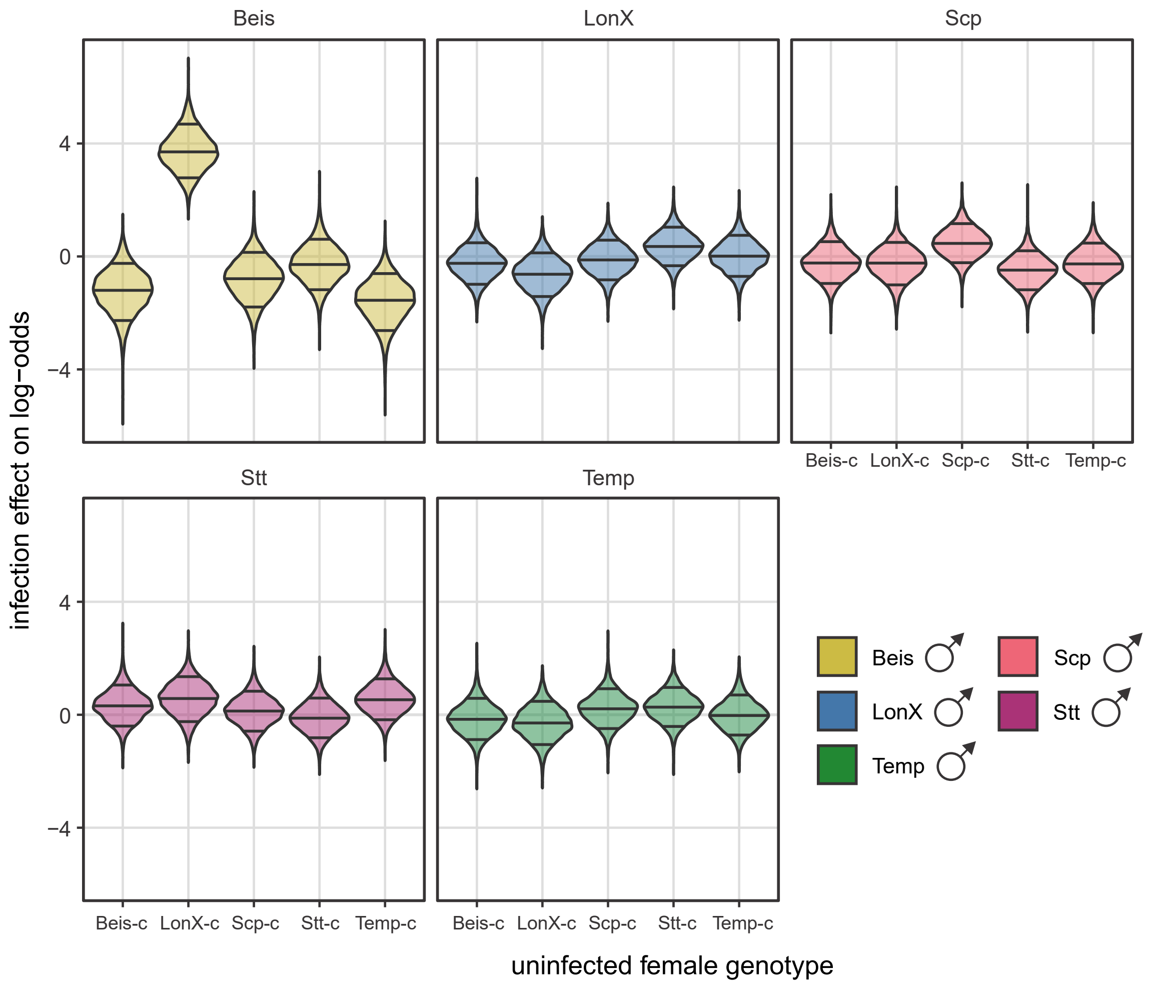


**Figure S4. Levels of interaction with the male infection state of intraspecific CI strength variation on the log-odds scale.** The panels are organized based on the male genotype. Male genotype is colour coded (see bottom right).

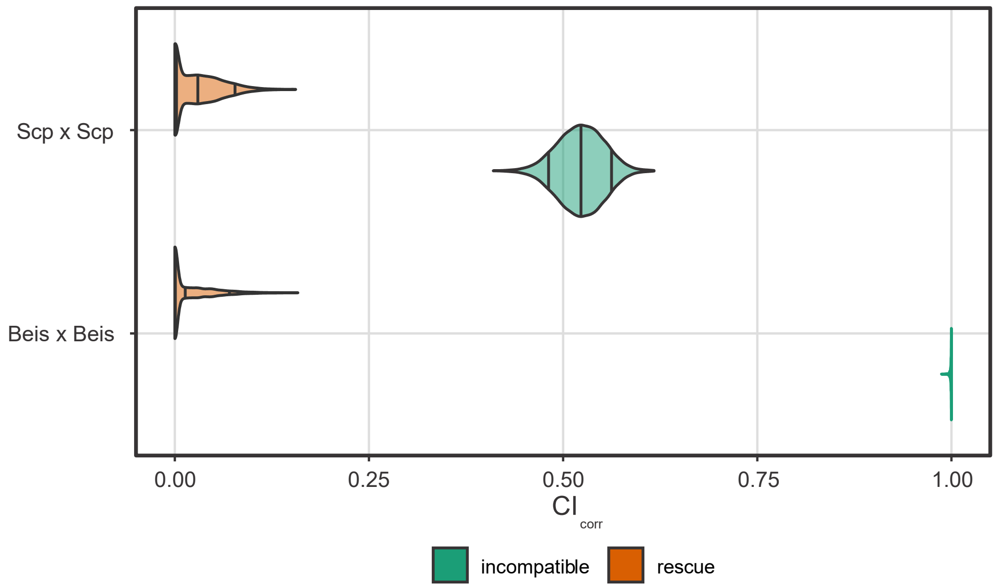


**Figure S5. Infected Beis-*w* and Scp-*w* females rescue CI.** CI strength was estimated using the $CI_{corr}$ index. Each violin plot represents the estimated average of $CI_{corr}$ for that cross. Rescue crosses were established using infected females, whereas incompatible crosses used uninfected females. $CI_{corr}$ estimates of the incompatible crosses are identical to those of Figure 1B. Distributions of estimates indicate 0.09, 0.50 and 0.91 percentiles.

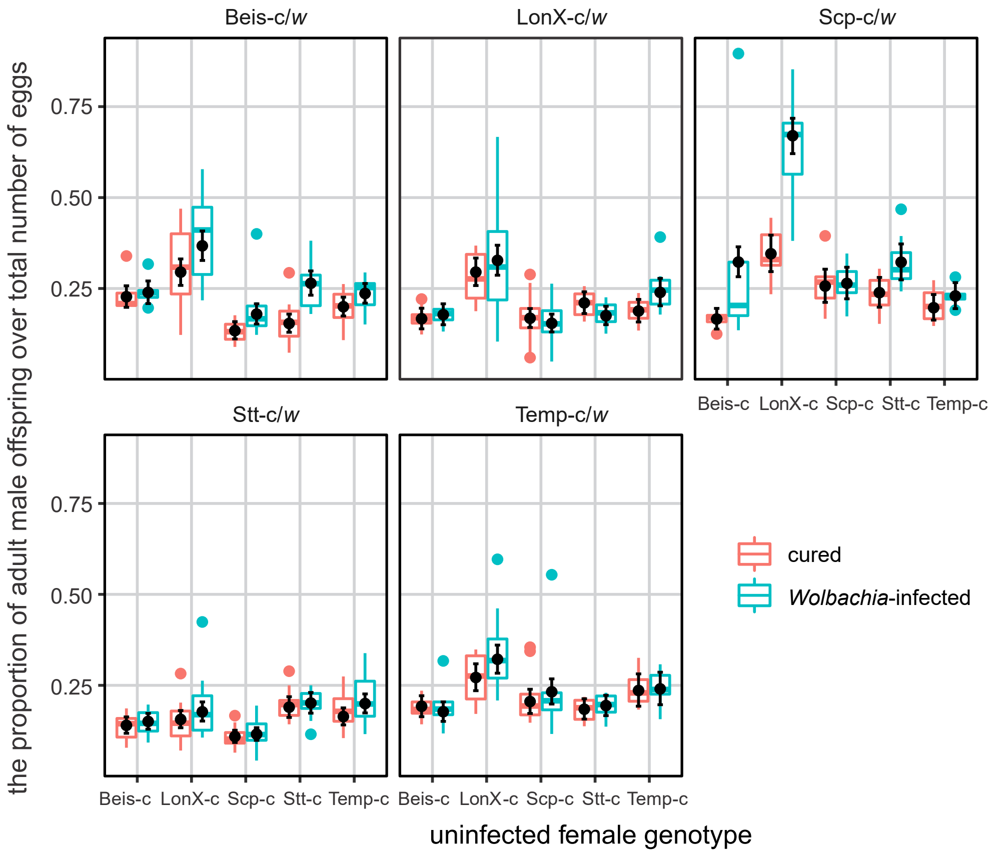


**Figure S6. Observed and estimated male proportion over total number of eggs across the intraspecific cross types.** The panels are organized based on the male genotype. The infection state of the males is colour coded (see bottom right).The boxplots depict the raw data whereas the dot and whiskers depict the mean model estimates and the 0.09-0.91 likelihood intervals respectively from the HMC model (model (8)) without the variable intercept for measuring day (electronic supplementary material).

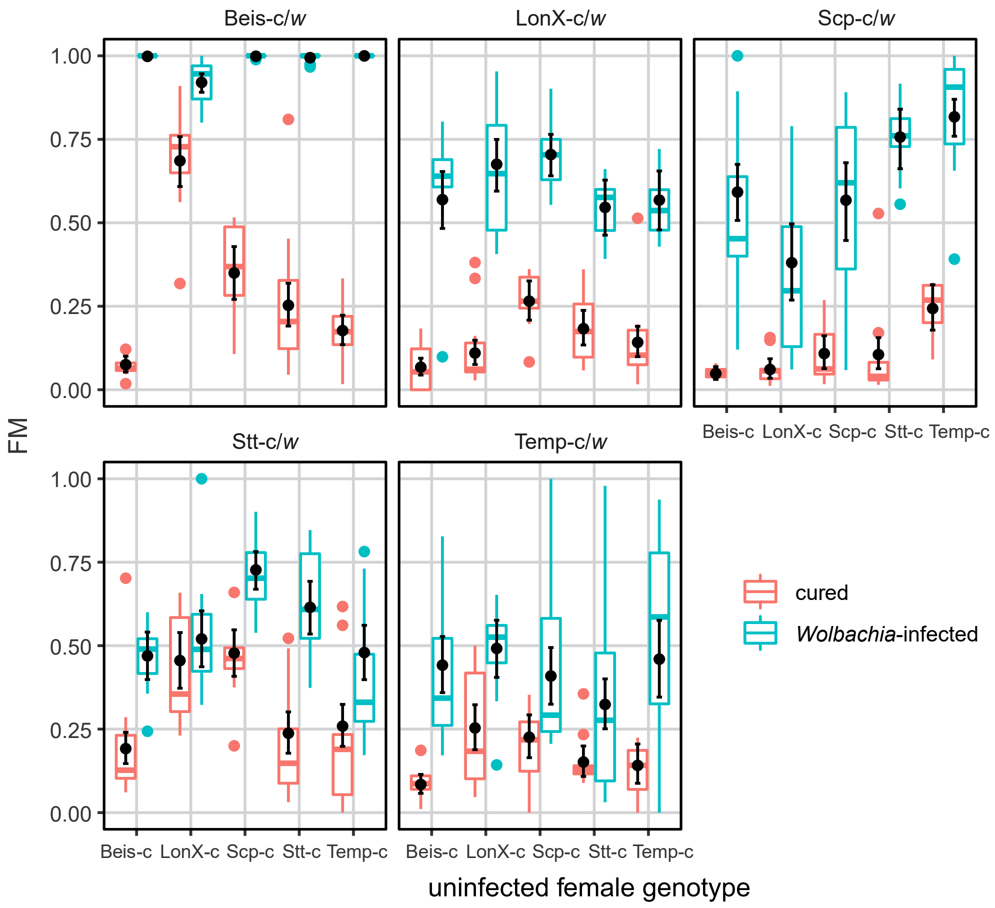


**Figure S7. Observed and estimated proportion of eggs that failed to generate adult mites over total number of eggs that did not generate adult males (FM) across the intraspecific cross types.** The panels are organized based on the male genotype. The infection state of the males is colour coded (see bottom right). The boxplots depict the raw data whereas the dot and whiskers depict the mean model estimates and the 0.09-0.91 likelihood intervals respectively from the HMC model (model (14)) without the variable intercept for measuring day (electronic supplementary material).

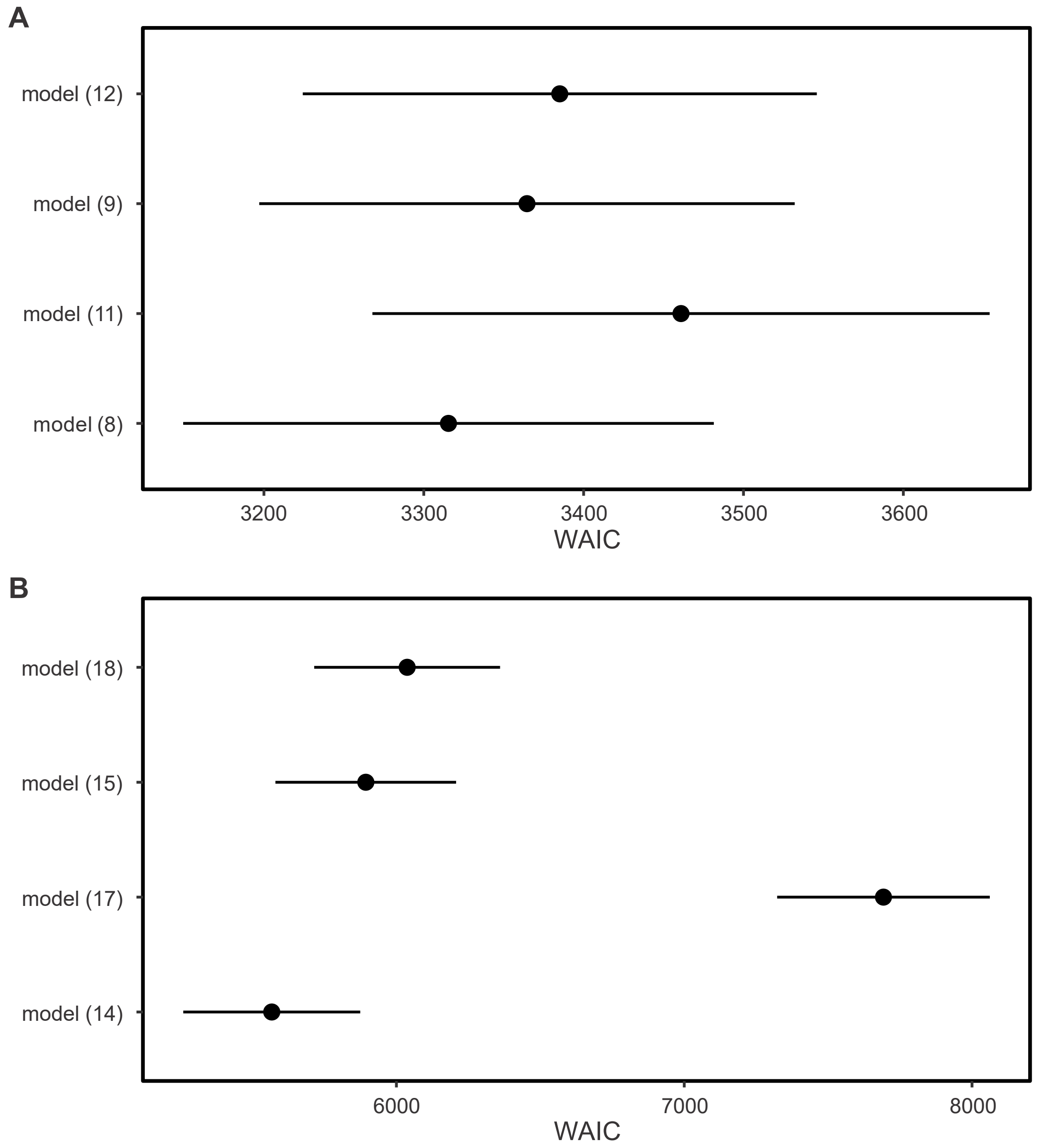


**Figure S8. Model comparisons for the analysis of the CI phenotypes in the intraspecific crosses.** For the MD-CI (A) and FM-CI (B) phenotypes, the WAIC scores and their standard errors are depicted for four models that were run to analyze intraspecific CI phenotype variation. For both panels, the metrics of four models are depicted; full models that include the effects of male and female genotype with their interaction as explanatory variables (model (8) and model (14)), models that incorporate the effects of male and female genotype without their interaction (model (9) and model (15)), models that incorporate the effects of female genotype (model (11) and model (17)) or male genotype (model (12) and model (18)). All models contained the effect of the *Wolbachia*-infection state of males, its interaction(s) with the other variable(s) and a random intercept for the days on which the cross types were initiated. Detailed model descriptions can be found in electronic supplementary material.

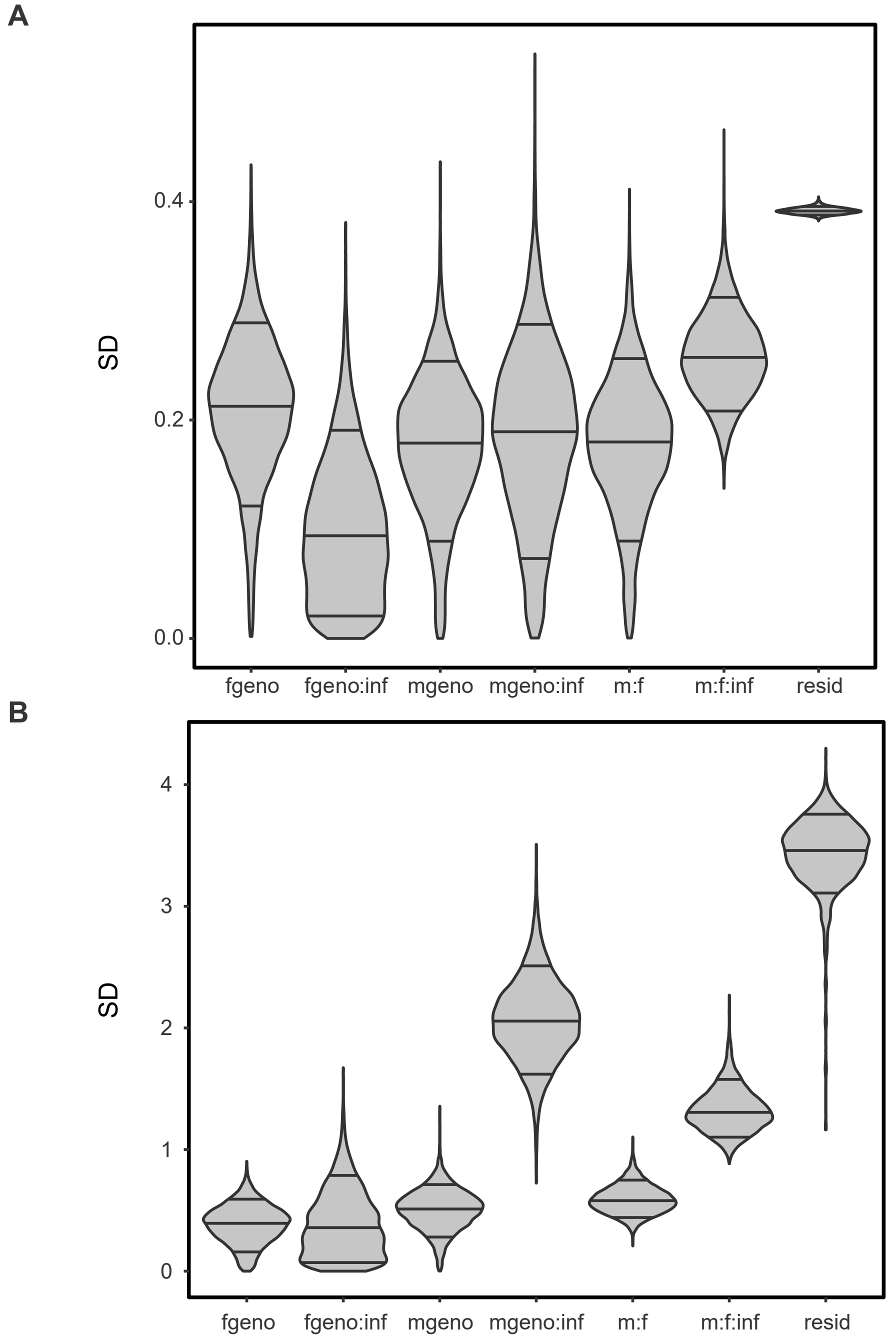


**Figure S9. Estimated finite-population standard deviation (SD) of coefficients of the levels of model (10) and (16) for the CI phenotypes to quantify the relative impact of the explanatory variables.** For the analysis of intraspecific CI phenotype variation, MD (A) and FM (B), the levels of the female genotype (fgeno), male genotype (mgeno), their interaction (f:m), and all the interactions with the *Wolbachia*-infection state in males (inf) were included. The standard deviations of the log-odds residuals of the model were included as references for the unexplained variance. The SD distributions show the 0.09, 0.5, and 0.91 percentiles. The detailed descriptions of model (10) and (16) can be found in electronic supplementary material.

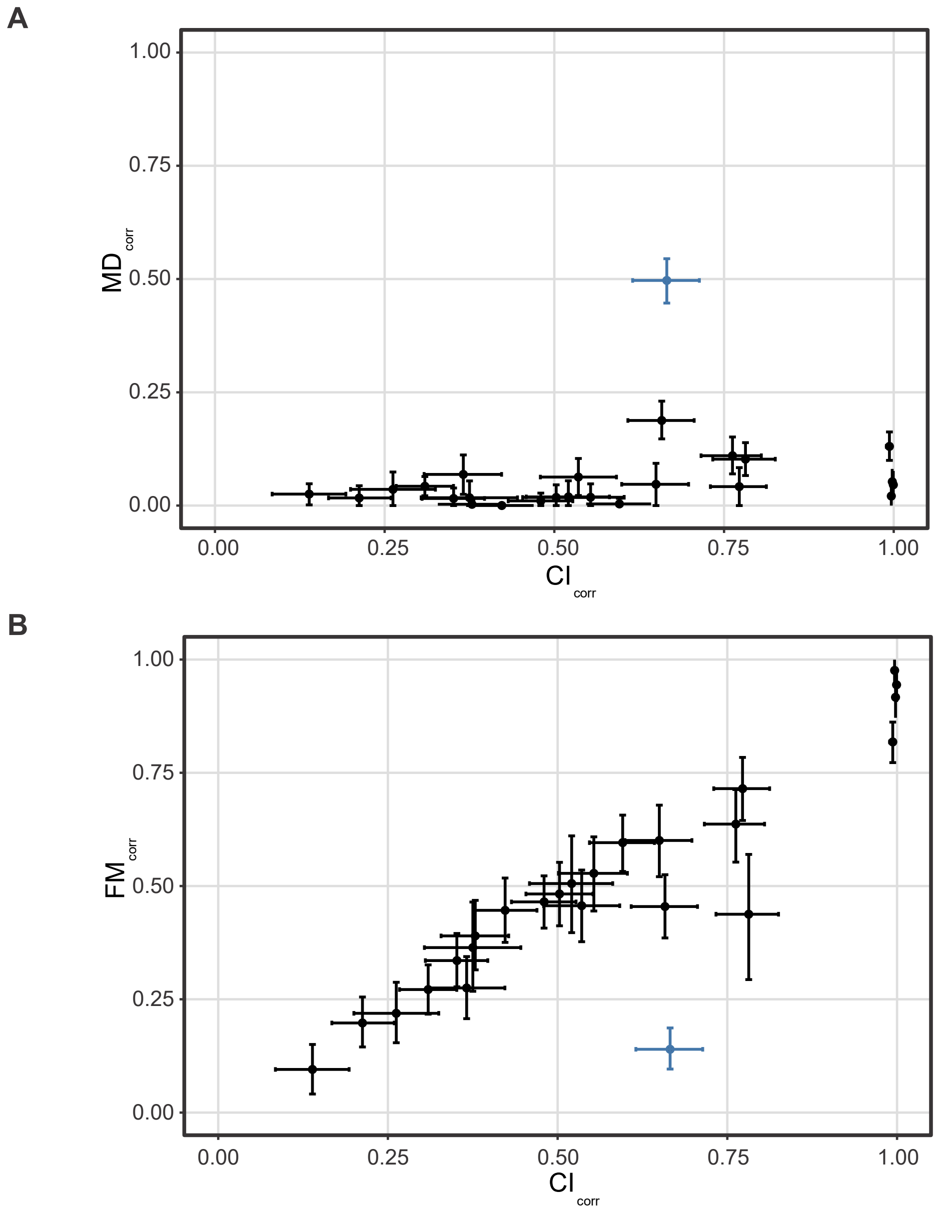


**Figure S10. Correlation of CI$_{corr}$ and MD$_{corr}$ (A) and FM$_{corr}$ (B) across the intraspecific cross types.** For both panels, the LonX-c (♀) x Scp (♂) cross is depicted in blue. Dots indicate estimated means and whiskers 0.09-0.91 likelihood intervals of the respective HMC model.

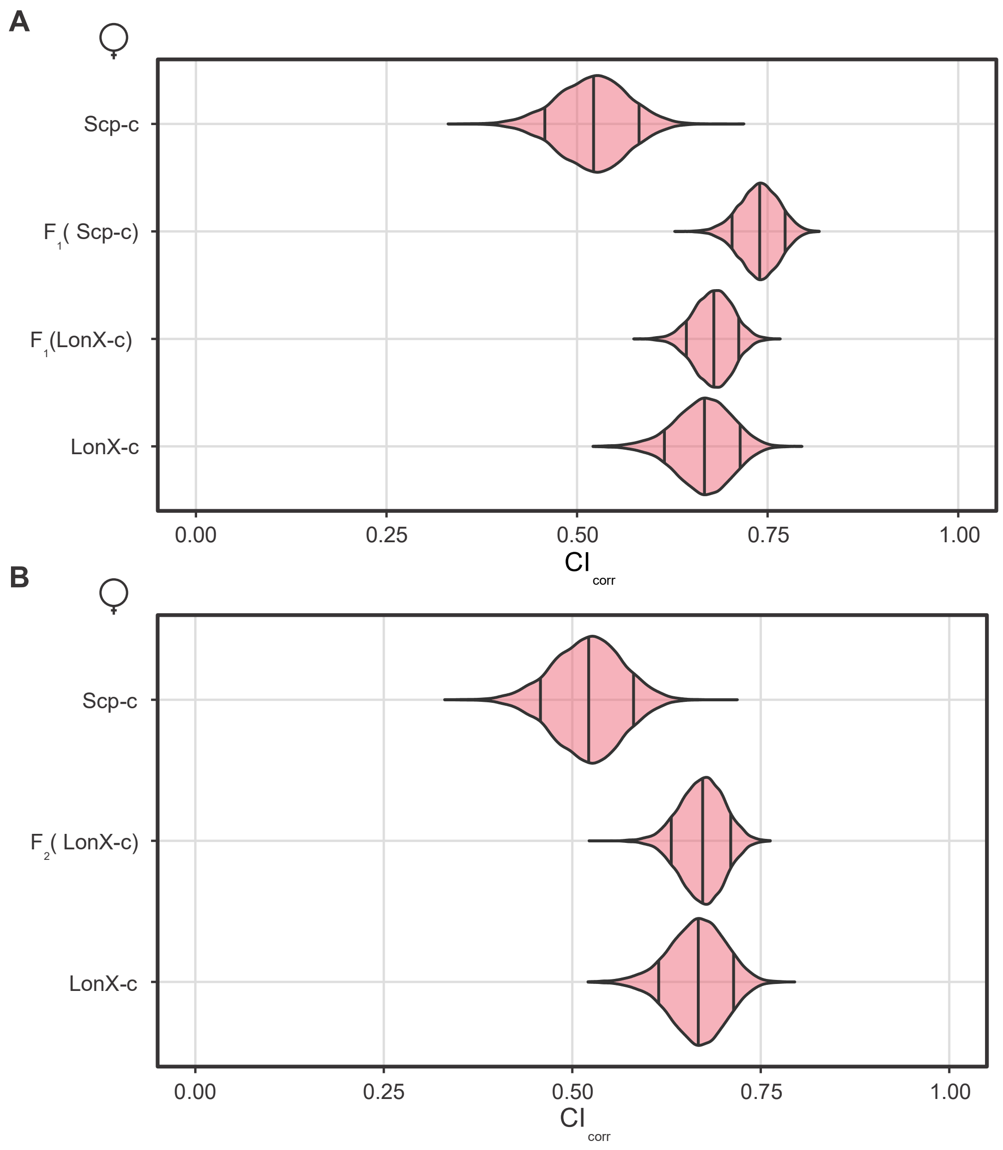


**Figure S11. CI strength remains stable for heterozygous $F_1$ females and recombinant $F_2$ females.** For the heterozygous $F_1$ and recombinant $F_2$ females, the genotype between brackets represents the original maternal genotype. All uninfected females were crossed to Scp-*w* and Scp-c males. $CI_{corr}$ estimates of LonX-c and Scp-c are identical to those of Figure 1B. Distributions of estimates indicate the 0.09, 0.50 and 0.91 percentiles.
