## Supplementary material for "Interacting host modifier systems control *Wolbachia*-induced cytoplasmic incompatibility in a haplodiploid mite": electronic supplementary material

### Supplementary materials and methods

#### Morphological classification of Bch

Following clearing in lactic acid (50%) for 24 h, Bch mites were mounted in Hoyer's medium. The specimens were examined using a Leica® DM LB 2 phase contrast microscope. Measurements were taken with live images using the software Amscope© suite (v. 3.7.7934) coupled with an AmScope® MU1803 camera.

#### Single mite DNA extraction

Individual mites were isolated and homogenized in 21 µl of PCR buffer (10 mM Tris-HCl, 100 mM NaCl, 1 mM EDTA, 2 mg/ml of proteinase K, with pH 8). Homogenates were incubated at 37°C for 30 min. Proteinase K was inactivated by incubating the mite samples at 95°C for 10 min. Each mite sample was diluted by adding 10 µl of sterile nuclease-free water.

#### Curing *T. urticae* of *Wolbachia* infection

Leaf discs of 16 $cm^2$ were dehydrated at 60°C for 1 min and soaked in a rifampicin solution (0.025 mg/ml). A cured line was created for each *Wolbachia*-infected line by transferring 40 larvae to a rifampicin-treated leaf disc and developing on rifampicin-treated leaf discs for two successive generations. Leaf discs were replaced every three days and rifampicin solutions were refreshed daily. Antibiotic curing of *Wolbachia* infection was confirmed by diagnostic PCR assays on 25 individual adult females and a pool of 100 adult females. Two adult females of the respective *Wolbachia*-infected sister line were processed in parallel and served as positive controls for the PCR assays. PCR conditions are described in Table S2.
