## Supplementary material for "Interacting host modifier systems control *Wolbachia*-induced cytoplasmic incompatibility in a haplodiploid mite": electronic supplementary material

### Supplementary results

Bch females belonged to the subgenus *Tetranychus* since they exhibited a diamond pattern on the dorsal striation. As Bch females bore four tactile setae proximal to the proximal duplex setae on tarsus I, they were further classified to the nineth species group as defined by Flechtmann & Knihinicki (2002) [1]. Here, identification to the species level is based on the size and shape of the male aedeagus (male external genital organ), more particularly its distal part called « knob ». Of the three observed Bch males, the shape of the aedeagus of one of them was similar to those observed in *Tetranychus urticae* [2] but, in the other two specimens, the combination between anterior and posterior projections of the knob gave the aedeagi a distinctly different appearance from that usually observed in *T. urticae*. The aedeagus of the three Bch males differed markedly from those of *T. urticae* in size. The knobs of the aedeagi observed were clearly larger since they varied from 2.9 to 3 µm versus ~2.5 µm in *T. urticae* [3]. To conclude, Ehara and Gotoh (1996) separated *T. urticae* morphologically from a sibling species, *Tetranychus pueraricola* based on the length of the aedeagus knob that varied around 2.5 and 2.1 µm for *T. urticae* and *T. pueraricola*, respectively [3]. Despite small differences in the knob length between the two species, the identity of *T. pueraricola* was later confirmed using molecular markers [4]. Because of the importance of the size of the knob of the aedeagus in the species identification within the genus *Tetranychus*, it is clear that the Bch specimens do not belong to the species *T. urticae*.
