## Supplementary material for "Interacting host modifier systems control *Wolbachia*-induced cytoplasmic incompatibility in a haplodiploid mite": electronic supplementary material

### Supplementary tables

### Table S1. Origins of the six *Tetranychus* genotypes

| Species | Name | Host plant | Sampling site and date |
|---|---|---|---|
| *T. urticae* | Beis | Solomon's seal | Brugge, Belgium (2020) |
| | LonX | Apple | Ontario, Canada (2000’s) |
| | Scp-*w* | Cucumber | Bredene, Belgium (2020) |
| | Stt | Rose | Gent, Belgium (2020) |
| | Temp | Frangipani | De Haan, Belgium (2020) |
| *Tetranychus* | Bch | Tomato | Brzeźnica, Poland (2020) |

### Table S2. PCR primers and annealing temperatures

| Species | Gene | Primer name | Primer sequence (5'-3') | Ann. temp |
|---|---|---|---|---|
| *Tetranychus* | *COI* | LCO1490 | GGTCAACAAATCATAAAGATATTGG | 48 °C |
| | | HCO2198 | TAAACTTCAGGGTGACCAAAAAATCA | |
| *Wolbachia* | *wsp* | *wsp*_81F | TGGTCCAATAAGTGATGAAGAAAC | 54 °C |
| | | *wsp*_691R | AAAAATTAAACGCTACTCCA | |
| | *gatB* | *gatB*_F1 | GAKTTAAAYCGYGCAGGBGTT | 54 °C |
| | | *gatB*_R1 | TGGYAAYTCRGGYAAAGATGA | |
| | *coxA* | *coxA*_F1 | TTGGRGCRATYAACTTTATAG | 54 °C |
| | | *coxA*_R1 | CTAAAGACTTTKACRCCAGT | |
| | *hcpA* | *hcpA*_F1 | GAAATARCAGTTGCTGCAAA | 54 °C |
| | | *hcpA*_R1 | GAAAGTYRAGCAAGYTCTG | |
| | *ftsZ* | *ftsZ*_F1 | ATYATGGARCATATAAARGATAG | 54 °C |
| | | *ftsZ*_R1 | TCRAGYAATGGATTRGATAT | |
| | *fbpA* | *fbpA*_F1 | GCTGCTCCRCTTGGYWTGAT | 55 °C |
| | | *fbpA*_R1 | CCRCCAGARAAAAYYACTATTC | |
| *Rickettsia* | *gtlA* | RICS741F | CATCCGGAGCTAATGGTTTTGC | 52 °C |
| | | RCIT1197R | CATTTCTTTCCATTGTGCCATC | |
| *Cardinium* | 16S rRNA | ChF | TACTGTAAGAATAAGCACCGGC | 52 °C |
| | | ChR | GTGGATCACTTAACGCTTTCG | |
| *Spiroplasma* | spacer region | SpitsJ04_F | GCCAGAAGTCAGTGTCCTAACCG | 52 °C |
| | | SpitsN55_R | ATTCCAAGGCATCCACCATACG | |

Thirty cycles were run for all PCR reactions. *Wolbachia* infection was tested using the primers that amplify a fragment of *wsp*. The diagnostic PCR assays for the detection of reproductive manipulators have been extensively tested on spider mites and other arthropods in previous studies.

**Table S3. *Wolbachia* maternal transmission in the infected near-isogenic lines**

| **Name** | ♀ | ♂ |
|---|---|---|
| Beis-*w* | 35 | 13 |
| LonX-*w* | 29 | 10 |
| Scp-*w* | 31 | 12 |
| Stt-*w* | 30 | 13 |
| Temp-*w* | 31 | 14 |

All adult females and males were infected with *Wolbachia,* indicating complete maternal transmission in the five infected near-isogenic lines.
